# The impact of obesity; key features of raised intracranial pressure and clinical sequalae in rodents

**DOI:** 10.1101/2021.08.05.455266

**Authors:** Connar Stanley James Westgate, Snorre Malm Hagen, Ida Marchen Egerod Israelsen, Steffen Hamann, Rigmor Højland Jensen, Sajedeh Eftekhari

## Abstract

Elevated intracranial pressure (ICP) is observed in many brain disorders. Obesity has been linked to ICP pathogenesis in disorders such as idiopathic intracranial pressure (IIH). We investigated the effect of diet induced obesity (DIO) on ICP and clinically relevant sequelae.

Rats were fed either a control or high fat diet. Following weight gain long term ICP, headache behavior, body composition and retinal outcome were examined. Post-hoc analysis of retinal histology and molecular analysis of choroid plexus and trigeminal ganglion (TG) were performed.

DIO rats demonstrated raised ICP by 55% which correlated with the abdominal fat percentage and increased non-respiratory slow waves, suggestive of altered cerebral compliance. Concurrently, DIO rats demonstrated a specific cephalic cutaneous allodynia which negatively correlated with the abdominal fat percentage. This sensitivity was associated with increased expression of headache markers in TG. Additionally, DIO rats had an *in vivo* increased retinal nerve fiber layer thickness associated with raised ICP with a subsequent post-hoc demonstration of neuroretinal degeneration.

This study demonstrates for the first time that DIO leads to raised ICP and subsequent clinically relevant symptom development. This novel model of non-traumatic raised ICP could expand the knowledge regarding disorders with elevated ICP such as IIH.

## Introduction

Elevation in intracranial pressure (ICP) is observed in a range of cerebral pathologies, such as traumatic brain injury (TBI), ischemic stroke, hydrocephalus and idiopathic intracranial hypertension (IIH). These disorders are associated with significant morbidity and are amongst the most debilitating, costly brain disorders ^1^. Currently, limited pharmacotherapies exist to treat patients with elevated ICP, as the mechanisms governing both normal cerebrospinal fluid (CSF) production, ICP regulation and their dysregulation in pathology remain unresolved. The molecular pathway by which water molecules cross the choroid plexus (CP) epithelial cells are not fully characterized, however the characterization of aquaporin water channels (AQPs) and water co-transporting proteins as the Na^+^-K^+^-2Cl^-^-1 cotransporter (NKCC1) have improved our understanding of this process ^2^.

The normal range of ICP in healthy adults is around −2 to 5 mmHg and is influenced by body positions, aging and diurnal variations ^3–5^. In recent years, studies have suggested that obesity may also be associated with ICP dynamics indicated by an association between elevated BMI and ICP ^6,7^. In support, a direct correlation between BMI, percentage body fat and CSF pressure has been demonstrated ^8^. Furthermore, obesity is the major risk factor for development of IIH, which is a metabolic disease of increased ICP without identifiable cause. The condition affects mainly women and 90% of the patients are obese ^9^. IIH has significant morbidity and the most common symptoms are chronic headache, cognitive impairment and impaired vision due to papilledema ^10,11^. The severity of papilledema and visual loss is associated with increasing BMI in IIH ^12,13^. Headache is also associated with other raised ICP disorders such as TBI and hydrocephalus. Obesity is also associated with migraine and is a risk factor for chronic headache ^14,15^.

The nature of IIH, with its atraumatic intracranial hypertension, chronic headache and impaired vision renders IIH an appealing model condition of elevated ICP that could provide valuable new knowledge regarding both regulation and dysregulation of ICP dynamics in pathology. However, no proper preclinical model for non-traumatic raised ICP or IIH exists. Our group developed the first rodent model mimicking aspects of obesity-induced raised ICP using Zucker rats (genetic model of obesity) ^16^. However, the increasing rate of obesity suggests that environmental and behavioral factors including dietary factors have been the major contributors to the obesity epidemic rather than rare genetic changes ^17^. There is no evidence that IIH is a genetic disease, as such the source of the disease is likely behavioral and environmental. Therefore, we have used DIO rats in this study which share many characteristics with the common form of human obesity. The aim was to explore the relationship between DIO, raised ICP, headache and structural morphology of the optic nerve head (ONH) and neuroretina using digital high-resolution and image-guided spectral domain optical coherence tomography (OCT). A novel and validated telemetric system to monitor ICP was used ^18^. In order to understand the molecular changes at CP and a headache relevant region, trigeminal ganglion (TG), molecular analysis was performed.

## Results

### Diet-induced obesity

Following weight gain, DIO rats selected for ICP surgery were 15% heavier (370±35.5 vs 316±22.9 g, t_(22)_=4.256, P=0.0003) with double the abdominal fat mass (49.4±11.4 vs 22.7±3.8 g, t_(17.5)_=8.351, P<0.0001), higher abdominal fat percentage (43.2±6.2 vs 27.9±2.3 %, t_(17.5)_=8.323 P<0.0001) and a marginally higher abdominal lean mass (65.4±15 vs 58.5±7.68 g, t_(22)_=2.211, P=0.03) than selected controls. There was no difference in fasting glucose (6.3±0.5 vs 5.6±0.8 mmol/L, t_(9)_=1.699, P=0.12).

### ICP recording

ICP telemeters were implanted in DIO and control rats in which ICP was continuously measured for 30 days. ICP was higher in DIO rats over the first 14 days (F_(1,17)_=12.00, P=0.003, Fig. 1A), where ICP on the day of surgery, day 0, correlated with abdominal fat percentage (r=0.54, P=0.016, Fig. 1B). On day 0, DIO rats had raised ICP relative to control rats (2.77±0.57 vs −0.17±0.73 mmHg, t_(17)_=3.202, P=0.0052, Fig. 1C, D). This raised ICP was associated with a right shift in the ICP pressure frequency histogram (Fig. 1E). On day three, a day not influenced by post-surgery drugs, ICP was more than double in the DIO rats (4.6±0.74 mmHg vs1.46±0.06, t_(16)_=3.199, P=0.0052, Fig. 1F,G) with a right shift in the ICP histogram (Fig. 1H). Similarly, on day 7 when the rats have fully recovered from surgery ^18^, DIO rats had a higher ICP (5.17±0.47 vs 3.53±0.55 mmHg, t_(16)_=2.268, P=0.03, Fig. 1I,J) with a right shift in the ICP histogram (Fig. 1K). From day 15 to 30 of recording there was no difference in ICP (F_(1,12)_=1.727, P=0.07, Fig. 1L), although due to technical issues not all days were recorded. In the final 7 days recording, DIO rats had a trend to higher ICP than the control rats (5.94±0.55 vs 4.32±0.6 mmHg, t_(16)_=1.907 P=0.07, Fig. 1M). This loss of significance in ICP could be mediated by weight loss. Although both groups lost weight as expected, DIO rats lost more weight before regaining weight to a lesser amount (−7.5±0.6% vs 0.6±1.7%, P=0.0004, Fig. 1N) ^18,19^.

**Figure 1.**
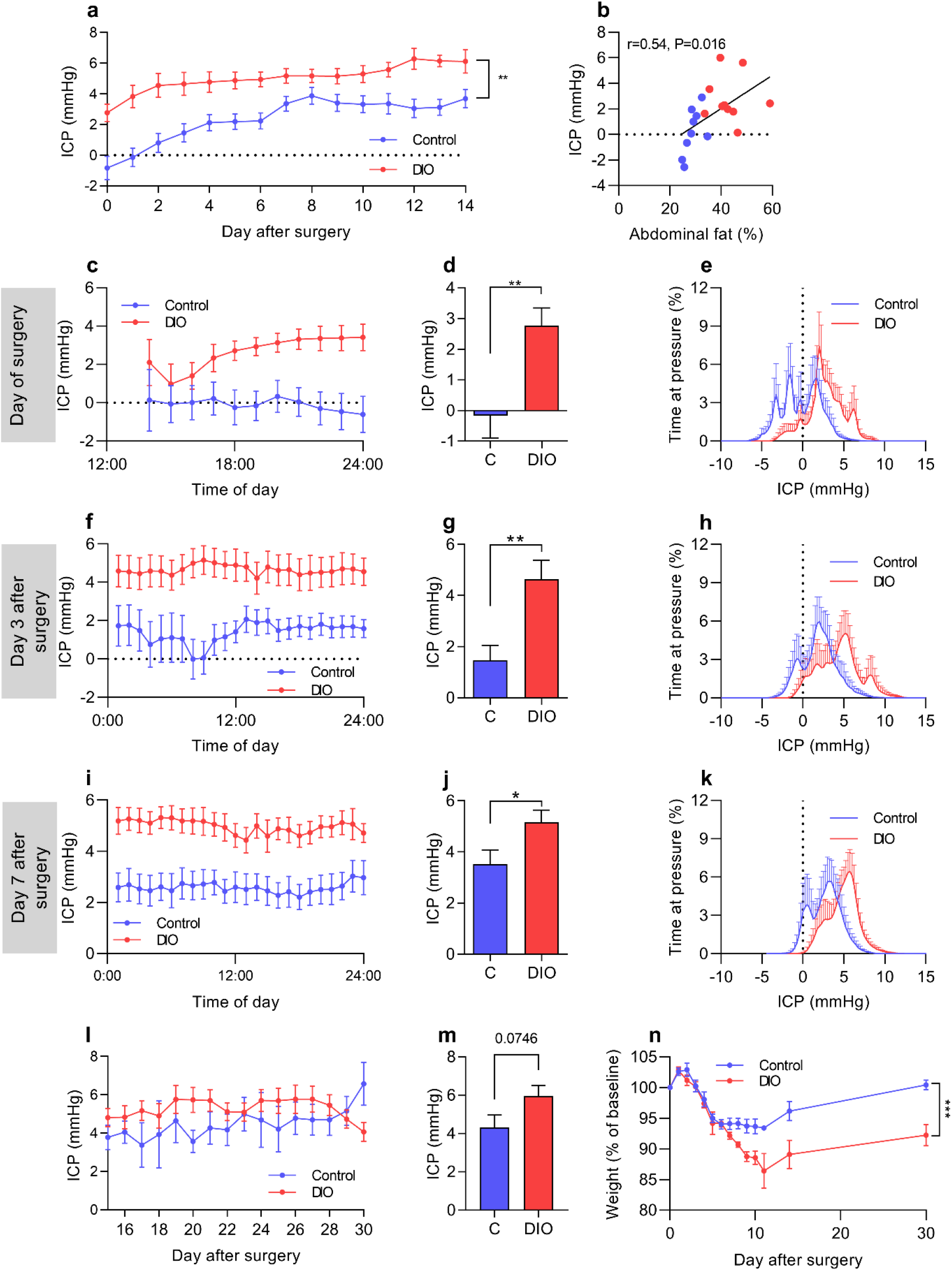
DIO increases ICP in female rats. Female Sprague-Dawley rats on control and very high fat diet implanted with ICP probes for 30 days. (**A**) ICP pressure trace over the 14 day ICP monitoring period. (**B**) Scatter graph depicting abdominal fat % vs ICP on day 0. (**C**) ICP pressure trace on day 0. (**D**) Mean ICP on the day of surgery. (**E**) Pressure frequency histogram from day of surgery (**F**) ICP pressure trace on day 3. (**G**) Mean ICP on day 3. (**H**) Pressure frequency histogram from day 3 (**I**) ICP pressure trace on day 7. (**J**) Mean ICP on day 7. (**K**) Pressure frequency histogram from day 7. (**L**) ICP trace from day 15 to 30, note that multiple data points are missing. (**M**) Mean ICP over the final 7 days of ICP recording. (**N**) Weight loss and regain following surgery. Control N=9, DIO N=10. Two-way repeated measures ANOVA for A,L and N. Pearson’s correlation for B and unpaired students T-tests for D, G, J and M. *=P<0.05, **=P<0.01. Data presented as mean±SEM.

To further analyze the ICP phenotype, ICP waveforms were assessed. On the day 0, there was no difference in any ICP frequencies, including those of slow non-respiratory ICP waves which are associated with pathological ICP features (0.29±0.07 vs 0.35±0.15 mmHg^2^, U=37, P=0.4, Fig. 2A, B). However, we identified a doubling in spectral power of ICP wavelengths at 0-0.25Hz (0.49±0.09 vs 0.20±0.08 mmHg^2^, U=16, P=0.03, Fig. 2C, D) on the third day after surgery. Similarly, there was a large increase in the spectral power of ICP wavelengths at 0-0.25Hz representing non-respiratory slow ICP waves (1.09±0.3 vs 0.24±0.09mmHg^2^, U=4, P=0.02, Fig. 2E, F) 7 days after surgery.

**Figure 2.**
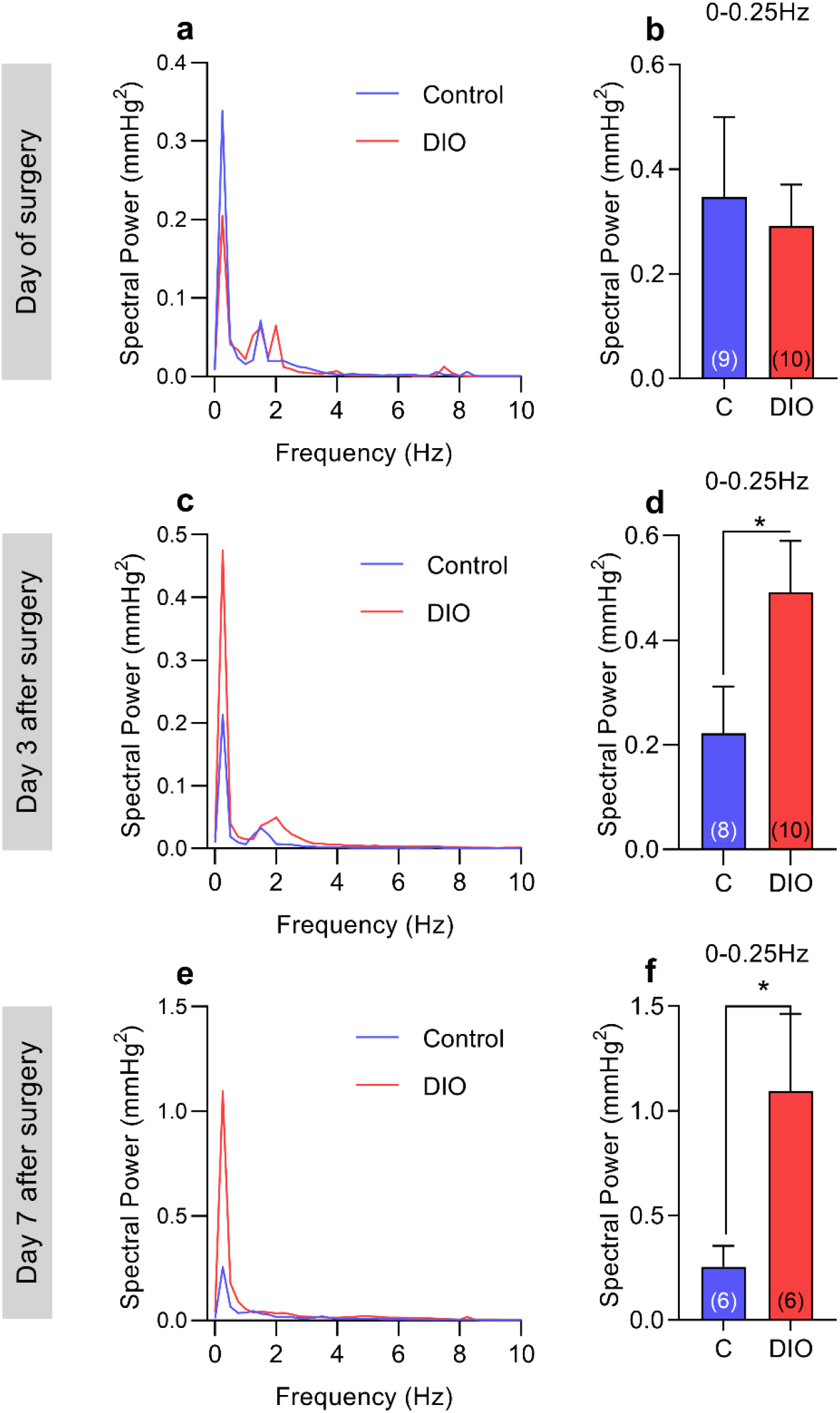
DIO alters ICP waveforms. Fast-Fourier transformation analysis of ICP waveforms between control and DIO rats. Power spectrum of ICP on day 0 (**A**) 3 days after surgery (**C**) and 7 days after surgery (**E**), condensed into 0.25Hz bins. Spectral power of slow ICP waves on day 0 (**B**), 3 days after surgery (**D**) and 7 days after surgery (**F**). N for each time point can be found in parentheses in the figure. Unpaired T-tests for B,D and F. *=P<0.05. Data presented as mean for A,C and E, mean±SD for B,D and F.

### Mechanical threshold, light sensitivity testing and expression in TG

All rats had mechanical thresholds assessed since the initiation of the diets. DIO rats had a lower threshold over the course of the diet (F_(1,149)_=7.318, P=0.0078, Fig. 3A), demonstrating cyclical periods of sensitivity. The hind paw data showed that no difference over the course of the diet (F_(1,231)_=1.142, P=0.28, Fig. 3B). DIO rats had lower periorbital (163.1±8.0 vs 213.8±5.1 g, t_(24)_=5.09 P<0.0001, Fig 3C) and hindpaw (211.6±6.7 vs 237.0±6.1 g, U=37, P=0.01, Fig. 3D) thresholds compared to controls, indicating prominent cephalic and mild peripheral mechanical cutaneous allodynia in DIO rats. Periorbital thresholds show a strong inverse correlation with abdominal fat percentage (r=-0.65, P=0.0005, Fig. 3E). ICP was not correlated with cephalic withdrawal thresholds on day 0 (r=-0.236, P=0.4). Light sensitivity was assessed in a light dark box and expressed as percentages of time spend in the light relative to time of the test. The control and DIO rats spend equal time in the light during the test (23.8 % ± 3.5 vs 24.92 % ± 2.9, t_(30)_=0.23, P=0.81, Fig. 3G).

**Figure 3.**
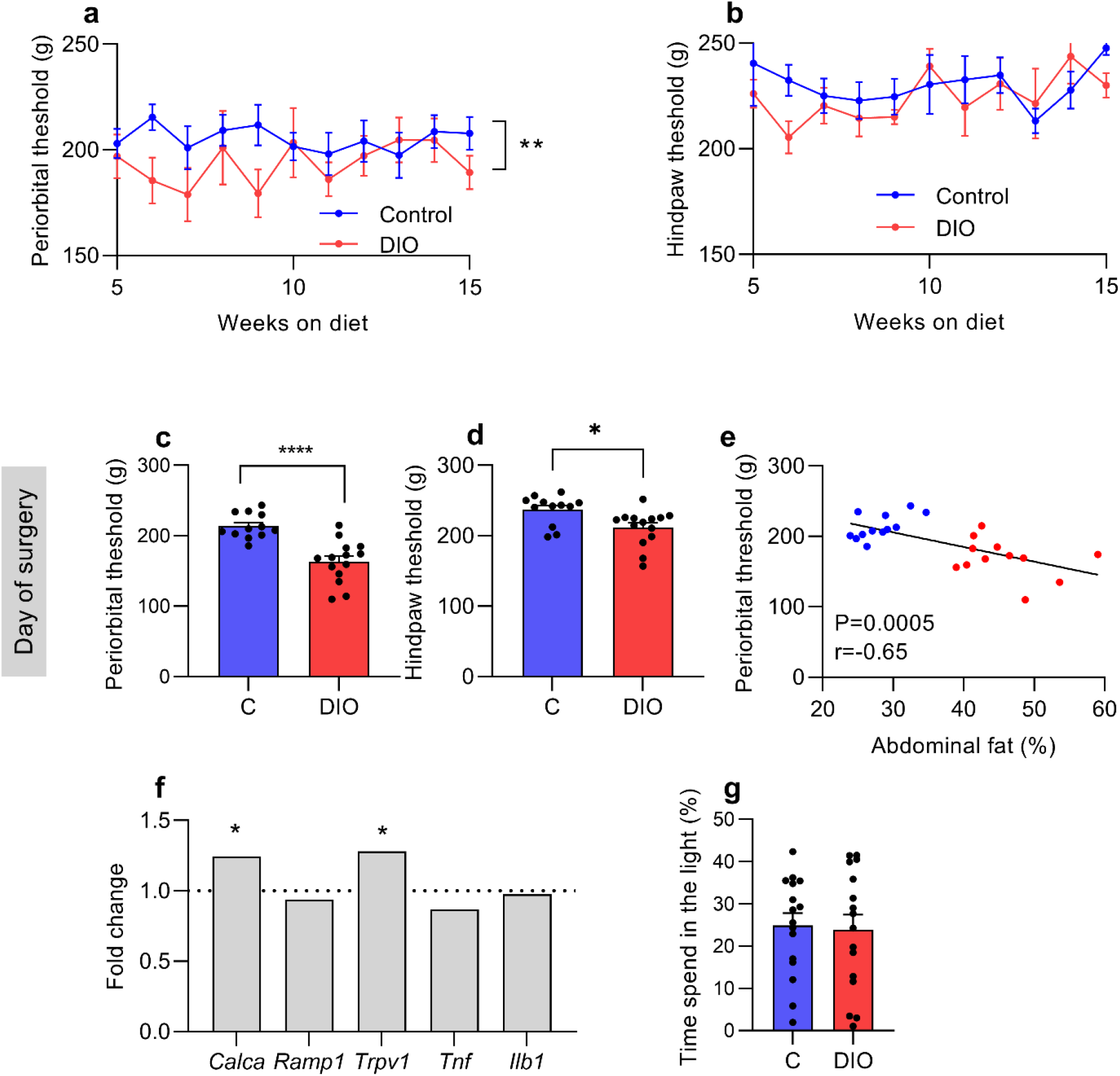
DIO induces cephalic allodynia. Peripheral and cephalic pain thresholds in female Sprague-Dawley rats on control and DIO rats. Longitudinal periorbital (**A**) and hindpaw (**B**) Von frey thresholds during weight gain. Periorbital (**C**) and hindpaw (**D**) Von frey testing on day 0. (**E**) Scatter graph of abdominal fat vs periorbital threshold on day 0, blue dots represent controls, red dots represent HFD rats. (**F**) Fold change in expression of *Calca, Ramp1, Trpv1, Tnf* and *Ilb1* in trigeminal ganglia of DIO rats, dotted line represents controls. (**G**) Photophobia testing of rats two weeks prior to surgery (N=16 control and DIO). Two-way repeated measure ANOVA for A and B. Unpaired students T-test for C and D. Pearson’s correlation for E. *=P<0.05, **P<0.01 and ****=P<0.0001. Control N=12 and DIO= 14. N=9 for TG RT-qPCR. Data presented as mean±SEM.

To further interrogate the headache phenotype, we assessed gene expression in TG on day 30 (Fig. 3F). In the DIO rats, there was a 1.24 fold increase in *Calca* (CGRPα) expression (2.9±0.05 vs 3.41±0.15, ΔCt, t_(15)_=2.544, P=0.02) and a 1.28 fold increase in *Trpv1* expression (5.94±0.0.06 vs 6.35±0.14 ΔCt, t_(15)_=2.551, P=0.02) compared to controls. No differences in expression in *Ramp1* (t_(15)_=0.68, P=0.58) or the pro-inflammatory genes *Tnf* (t_(15)_=0.98, P=0.78) and *Il1b* (t_(15)_=1.279, P=0.48) were found.

### Evaluation of ODP, OCT retinal layer segmentation and IOP

Raised ICP is associated with retinal abnormalities and potentially retinal degeneration. Given this, we assessed the retinal anatomy. The ODPs in both groups appeared normal without any notable pathology or any signs of acute papilledema (Fig. 4B). The OCT (Fig. 4A) scans showed no significant thickness difference between right and left eye in any of the retinal layers (Table 1). No significant thickness difference between DIO and control animals was found for TR, GCC, IRL and ORL (Table 1). RNFL was significantly thicker in the DIO (28.81±0.61 vs 24.85±1.093 μm, t_(22)_=3.390, P=0.0026, Fig. 4C) compared to control animals. Positive correlations were observed between RNFL thickness and ICP (r=0.639, P=0.0058, Fig. 4D), and weight (r=0.56, P=0.0048) and DEXA abdominal fat percentage (r=0.469, P=0.02, Fig. 4E). No significant difference in intraocular pressure was found (15.73±0.872 vs.13.72±0.760 mmHg, t_(17)_=1.714, P=0.104).

**Figure 4.**
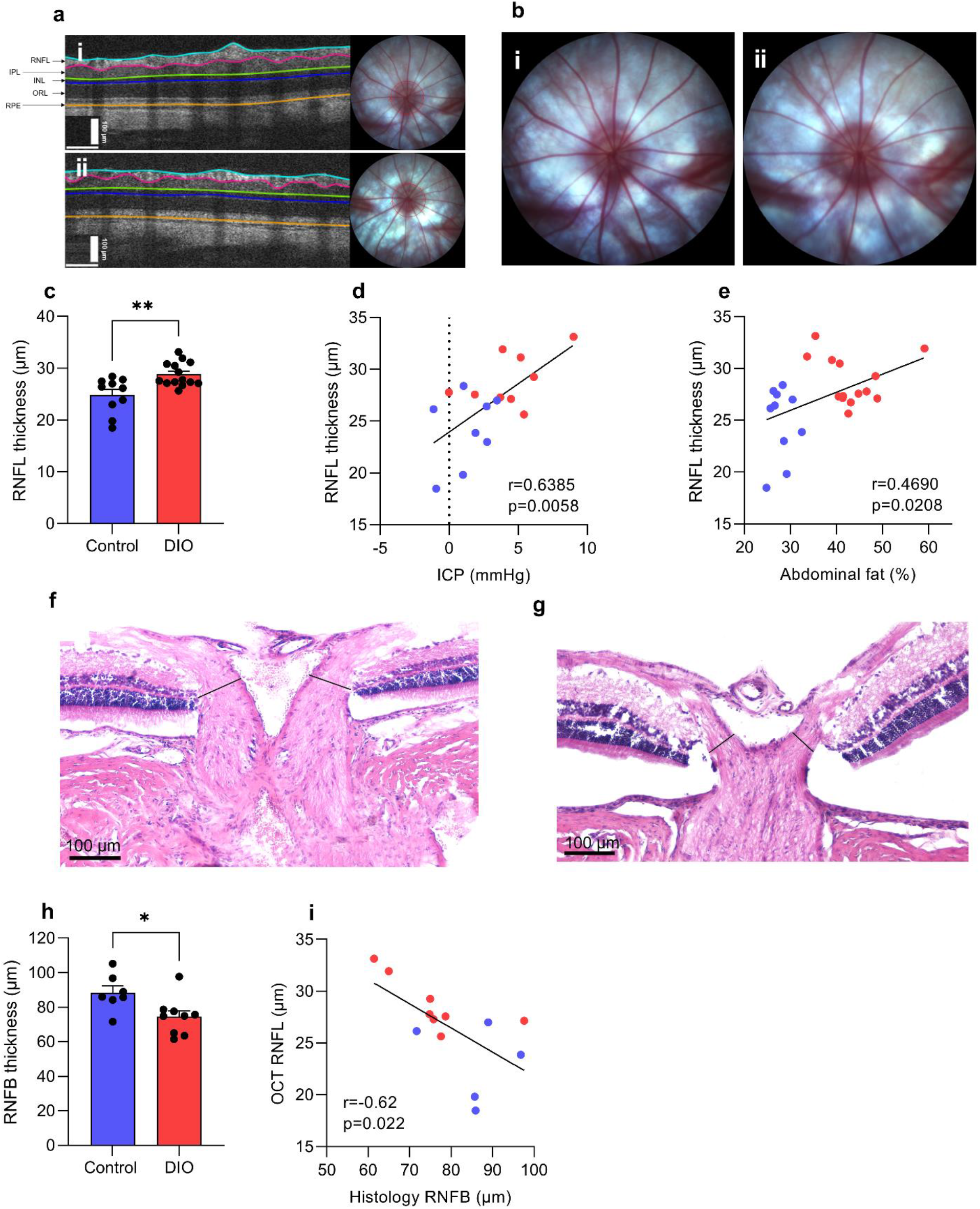
DIO and non-traumatic increased ICP alter neuroretinal structures. (**A**) Horizontally aligned OCT B-scans covering the peripapillary part circumferentially to the optic disc with manually adjusted segmentation lines in DIO (i) and control (ii) rats. (**B**) Optic disk photo in a DIO (**i**) and control animal (**ii**). (**C**) RNFL thickness between DIO and control. (**D**) Scatter graph showing a association between RNFL thickness and mean ICP on day 3. (**e**) Scatter graph showing association between RNFL thickness and abdominal fat-%. RNFB thickness in the prelaminar region of the ONH in a control animal (**F**) and a DIO animal (**G)** showing measure points (black solid lines). (**H**) Thickness of the RNFB. (**I**) Scatter graph showing correlation between OCT RNFL and histological RNFB. RNFL= retinal nerve fiber layer, IPL=inner plexiform layer, INL=inner nucleus layer, ORL=outer retinal layers (including outer plexiform layer, outer nucleus layer, inner segments and outer segments) and RPE=retinal pigment epithelium. Data in **C** and **H** are presented as mean±SEM. Unpaired T-test for C and H, Pearson’s correlation for D, E and I. *=P<0.05, **=P<0.01

**Table 1.** Retinal layer thickness. Table summarizing the thickness of the retinal layers at OCT prior to implantation surgery. N in parentheses. Data are presented as mean±SEM. **=P<0.01.

| OCT | DIO (14) |  |  | Control diet (10) |  |  |
| --- | --- | --- | --- | --- | --- | --- |
| Mean thickness (μm) | Both eyes | Right eye | Left eye | Both eyes | Right eye | Left eye |
| Total retinal | 172.85±2.20 | 172.61±2.49 | 173.09±2.30 | 169.56±2.76 | 169.30±2.62 | 169.82±3.59 |
| Retinal nerve fiber layer | 28.82±0.61** | 28.52±0.74 | 29.12±0.75 | 24.85±1.09 | 25.33±1.06 | 24.37±1.35 |
| Ganglion Cell complex | 65.64±1.06 | 66.14±1.22 | 65.15±1.15 | 65.00±1.34 | 64.69±1.59 | 65.32±1.31 |
| Inner retinal layers | 81.74±1.40 | 81.76±1.59 | 81.72±1.42 | 82.14±1.72 | 82.57±1.72 | 81.70±1.95 |
| Outer retinal layers | 91.11±1.40 | 90.84±1.78 | 91.38±1.39 | 87.41±1.62 | 86.73±1.74 | 88.10±2.64 |

To investigate the structural outcome of the retinal swelling, we investigated the thickness of the retinal nerve fiber bundles (RNFB) (Fig 4F and G). Here, we demonstrate that DIO animals had thinner RNFB (74.3±3.6 vs 88.3±3.9 μm, t_(14)_=2.590, P=0.0214, Fig. 4H) compared to control animals. Furthermore, we demonstrate a negative correlation between the thickness of the RNFL at OCT scan and the thickness of the RNFB (r=-0.62, P=0.022, Fig. 4I).

### Expression at CP

To investigate the molecular origin of the raised ICP, we assessed the expression of genes associated with CSF secretion at the CP. In CP, mRNA expression of *Nkcc1* in the DIO rats displayed a 1.11-fold change (−9.56±0.26 vs −9.40±0.19 ΔCt, t_(10)_=0.48, P=0.6) and a 1.45-fold change of *Aqp1* (−9.56±0.26 vs −9.40±0.19 ΔCt, t_(13)_=0.88, P=0.38) (Fig. 5A). The same trend was also observed by immunoblotting (Fig. 5B). Total Aqp1 was displayed a 2.8-fold increase in expression (5.3±1.5 vs 2.4±0.5, AU, t_(9.907)_=2.223) P=0.05, Fig. 5C), where by the proportion of glycosylated Aqp1 to total expression trended to increase in DIO rats (0.59±0.06 vs 0.46±0.03 %, t_(15)_=1.847 P=0.084, Fig. 5D). The protein expression of Nkcc1 was also higher by 1.28-fold (4.7±0.92 vs 3.4±0.78 AU, t_(15)_=0.79 P=0.43) (Fig. 5E).

**Figure 5.**
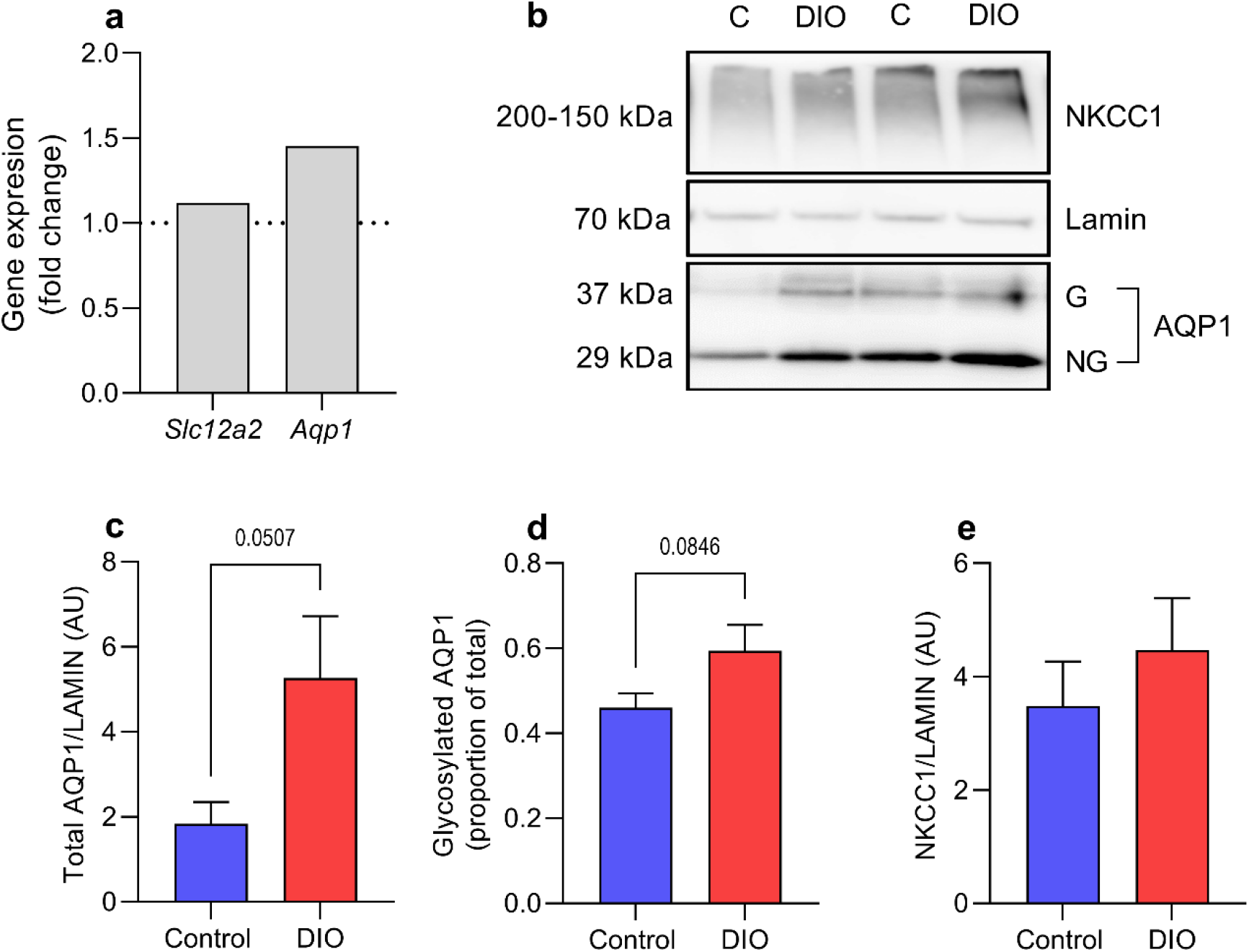
Expression of AQP1 and NKCC1 at CP. Expression of NKCC1 and AQP1 at CP in female Sprague-Dawley rats on control and HFD after 30 days implant with telemetric probes. Fold change of mRNA expression of NKCC1 and AQP1 in CP (**A**). Representative western blots (**B**). Protein expression of AQP1 (**C**), proportion of glycosylated AQP1 (**D**) and NKCC1 protein expression**(E**).Dotted line represent mean for controls in A. Unpaired students t-test is done for A, C, D and E. Data presented as geometric mean for A and mean±SEM for C, D and E. N=9.

## Discussion

Obesity is a global health issue, affecting over half a billion individuals with multiple adverse consequences ^20^. In recent years it has also been suggested that obesity could cause elevation in ICP ^8,16^. Elevated ICP is observed with several neurological disorders and if not treated, it is very likely to cause severe morbidity and even death if ICP rises too high. Therapeutic approaches for clinical management of elevated ICP are virtually absent, which may be due fact that the molecular mechanisms causing raised ICP remain unidentified. Furthermore, no suitable animal model for non-traumatic raised ICP exists. Only few animal models with elevated ICP exist and they all are associated with the induction of severe cerebral traumas. Thereby, in the current study we have investigated the impact of DIO on ICP and also developed an animal model for non-traumatic raised ICP including relevant symptom development for the first time. Here we show that DIO rats develop raised ICP, headache related hypersensitivity and neuroretinal changes.

IIH is a challenging disease with increased ICP of unknown etiology and is also closely linked to obesity. The incidence is rapidly increasing in the wake of the global obesity epidemic ^9^. Importantly, a recent study demonstrated that in non-IIH patients, BMI and percentage body fat both positively correlated with CSF opening pressure, although to a lesser degree than seen in IIH ^8^. In support, we have recently shown that with normal weight gain in rats, there was a trend to an increase in ICP, supporting the idea of obesity has an impact on ICP ^18^. In the current study, raised ICP was found in the DIO group, where the ICP was raised by 55% and associated with increased abdominal obesity. This study is congruent with our previous study in which it was shown that genetic obesity in rats caused 40% higher ICP ^16^. As in our previous study using the same telemetric system for ICP monitoring in normal rats, the resting ICP value for the rats at the day of surgery increased steadily on the following days and stabilized around day 7 ^18^. Previously, we demonstrated that ICP dynamics are influenced by recovery from surgery and the use of anesthesia alters the ICP waveforms ^21^. Interestingly, it was found that the DIO rats had increased non-respiratory slow waves after recovery from surgery, potentially indicating an altered cerebral compliance.

Additionally, the present study mimics human studies where it has been demonstrated in both IIH patients and other neurological patients that excess adiposity is associated raised ICP ^8,22^. The correlation between abdominal obesity and ICP in our model suggests that factors associated with adipose mass may affect ICP. No adipose derived factors in humans have been explicitly demonstrated to raise ICP. However, in both lean and obese female rats, exogenous glucocorticoids (GCs) hormones which are known to be increased in obesity, have been demonstrated to increase CSF secretion without altering CSF drainage suggesting that GC have the capacity to alter CSF dynamics ^23^. Whether the observed increase in ICP in the obese animals is due to obesity, endocrine abnormalities or other neurochemical disturbances caused by obesity is yet unknown. A limitation of the current study is not performing ICP recording during obesogenesis, thus preventing the observation of the temporal development of raised ICP. Although the use of female rats makes this work relevant to IIH and more generally females, further studies assessing the effect of DIO on male rats will help complete our understanding.

It has been demonstrated that female obese rats have increased CSF secretion ^23^. In the current non-traumatic raised ICP model, there was a trend to increased mRNA expression of *Aqp1* and *Slc12a2*. This trend was also observed in protein expression, where there was tendency to increased ratio of glycosylated AQP1 to total AQP1. Due to a slower weight gain after surgery in the DIO rats, the weight difference at 30 days was decreased to 3.4%. This may have affected the outcome in the molecular analysis and therefore requires further investigations. We have previously shown that obese Zucker rats with elevated ICP had increased expression of AQP1 at CP ^16^. Furthermore, AQP1 knockouts had 56% lower ICP in comparison with wild-type mice ^24^. All together this suggests that the obesity-induced ICP may modulate CSF dynamics by altering the expression of the water channels.

Headache is a frequent symptom in pathologies with raised ICP such as TBI, hydrocephalus and IIH ^25^. Headache is present in around 75-94% of IIH patients at diagnosis ^26–28^. In the present study, we show that DIO rats had cephalic cutaneous mechanical allodynia but not peripheral, where the severity was closely associated with higher levels of abdominal obesity. The altered pain thresholds were cyclical from 5 weeks on diet and until day of surgery. Cephalic cutaneous allodynia is considered to be a measurement of central sensitization to pain and marker of progression to migraine-like chronification in humans ^29^. The present study adds novel data to the few experimental studies suggesting that obesity increases headache like behavior in rodents, where DIO increased facial pain behavior, and in contrast to our data, photophobic behavior in mice ^30,31^. We did not find any difference in the light sensitivity between the DIO and control animals suggesting that these animals are not photophobic. Such difference could be accounted for different species between studies or use of fewer animals. Importantly, our data mirrors the human data, where obesity is associated with increased incidence of headache, and increased abdominal obesity confers both an increased risk of migraine diagnosis and increases cutaneous allodynia severity ^29,32^. No link between ICP and cephalic thresholds was found, mimicking the clinical features of IIH, where ICP did not correlate with the reported headache severity ^26^. In support to this, two-thirds of patients with IIH continue to experience disabling headaches after the other manifestations of the disorder including the ICP had normalized ^26^.

CGRP plays a key role in headache disorders and inhibition of CGRP signaling has demonstrated clinical efficacy as migraine therapy ^33^. Furthermore, increasing attention has been given to the capsaicin-gated channel transient receptor potential vanilloid subfamily member 1 (TRPV1), suggested to play a role in migraine as well as the sensitization phenomena related to it ^34^. For the first time, this study shows increased expression of CGRP and TRPV1 in a model of non-traumatic raised ICP, suggesting that these molecules may be involved in the headache related behavior observed in this model. Within the TG, CGRP is expressed in small-sized neurons, and the c-fibers in the dura mater and at the brain stem level, where c-fibers are nociceptive^35–37^ This suggests that obesity or raised ICP sensitizes the TG to pain. TRPV1, a thermoreceptor, is expressed in small to medium size nerve fibers and neurons in the TG, i.e nociceptive neurons and often colocalizes with CGRP. TRPV1 activation causes CGRP release from TG neurons^38,39^. Our demonstration of increased *Trpv1* expression is corroborated by previous *in vivo* data demonstrating that cultured TG neurons from obese mice are more sensitive to the TRPV1 agonist capsaicin^40^. TRPV1 expression is modulated by inflammatory mediators, given that the TG is outside the blood brain barrier, this suggests that the circulating inflammatory milieu in obesity could be causative in the upregulation of TRPV1, given that our qPCR data suggests no increases in pro-inflammatory gene expression in the TG^21,41^. The direct cause of the molecular and therefore the cutaneous allodynia remains uncertain. The raised ICP through direct mechanical stimulation of the dura mater and TG could increase sensitivity. Additionally, obesity related factors could be also involved. Further work to elucidate the underpinnings of the cutaneous allodynia is required.

Severe obesity is found to be related to retinal neurodegeneration ^42^. In this study, the OCT data showed thickening of the peripapillary RNFL in DIO rats with a significant positive correlation to ICP. Across both the DIO and the control animals, there was also a clear correlation between RNFL thickening and the total body weight and as well the abdominal fat percentage. This identifies abdominal obesity as a potentially disease aggravating factor. We suggest this *in vivo* RNFL thickening observed with OCT may be due to swelling of the retinal ganglion cell axons due to axoplasmic flow stasis. This initial RNFL swelling is also present in other models with varying optic nerve or disc damage^43,44^. By histology, we showed that after a prolonged period of DIO and elevated ICP there was a significant thinning of RNFBs entering the optic disc. This is in accordance with the clinical setting where obesity is found to correlate with both raised ICP and RNFL thinning ^42,45^. Furthermore, we demonstrate that the degree of RNFL swelling at OCT is negatively associated with RNFB thickness at histology, i.e the degree of swelling correlates with the degree of neurodegeneration. To our knowledge this is the first animal model to demonstrate a link between obesity induced non-traumatic ICP elevation and retinal nerve fiber degeneration. The link between ICP and neuroretinal degeneration has previously been demonstrated in mouse models of sustained and robust elevated ICP which caused fundoscopic visible papilledema with neuroretinal and optic nerve degeneration ^46,47^. However, in the present study no signs of acute papilledema were detected, which may be explained by the moderate ICP increase that occurs in this non-traumatic model. Even in a rat model with markedly increased ICP > 15 mmHg only modest signs of acute papilledema are detected. Future studies using electrophysiological techniques, OCT, histology and ICP-recordings combined are needed to further understand the retinal function and how it varies with weight changes.

Taken together, this study reveals for the first time the impact of DIO on ICP with clinically relevant symptom development. These findings contribute to increasing the understanding of the impact of obesity on brain health and the neurobiology of raised ICP disorders, especially IIH. This non-traumatic raised ICP model provides the tools that can be utilized in the future to evaluate underlying mechanisms and potential ICP reducing agents to determine their efficacy.

## Material and Methods

### Animals and diets

46 female Sprague-Dawley (SD) rats (Taconic, Denmark) up to 12 weeks of age were started on the diet. At day of arrival they were placed on either 60% high fat diet (HFD) or matched control diet (CD) with 10% fat for 15 weeks (Fig. 6) (Research Diets, D12450J and D12492) housed in the animal facility at Glostrup Research Institute, Denmark. Rats were left a week to acclimatize prior to all procedures. Food consumption and body weight were monitored weekly. Weight-based selection was made in the end of the diet time for ICP implantation and eye examination (Fig.6). In the HFD group (N=22), the top rats by weight were selected for surgery, whereas in the control diet group (N=24), the bottom rats by weight were selected for surgery. The study was approved by the Institutional Animal Care and Use Committee (Glostrup Research Institute) and the procedures were carried out in accidence to Danish Law. Licenses granted by the Danish Animal Experiments Inspectorate ( license numbers 2014-15-0201-00256, 2019-15-0201-00365). All methods and data are reported according to ARRIVE guidelines.

**Figure 6.**
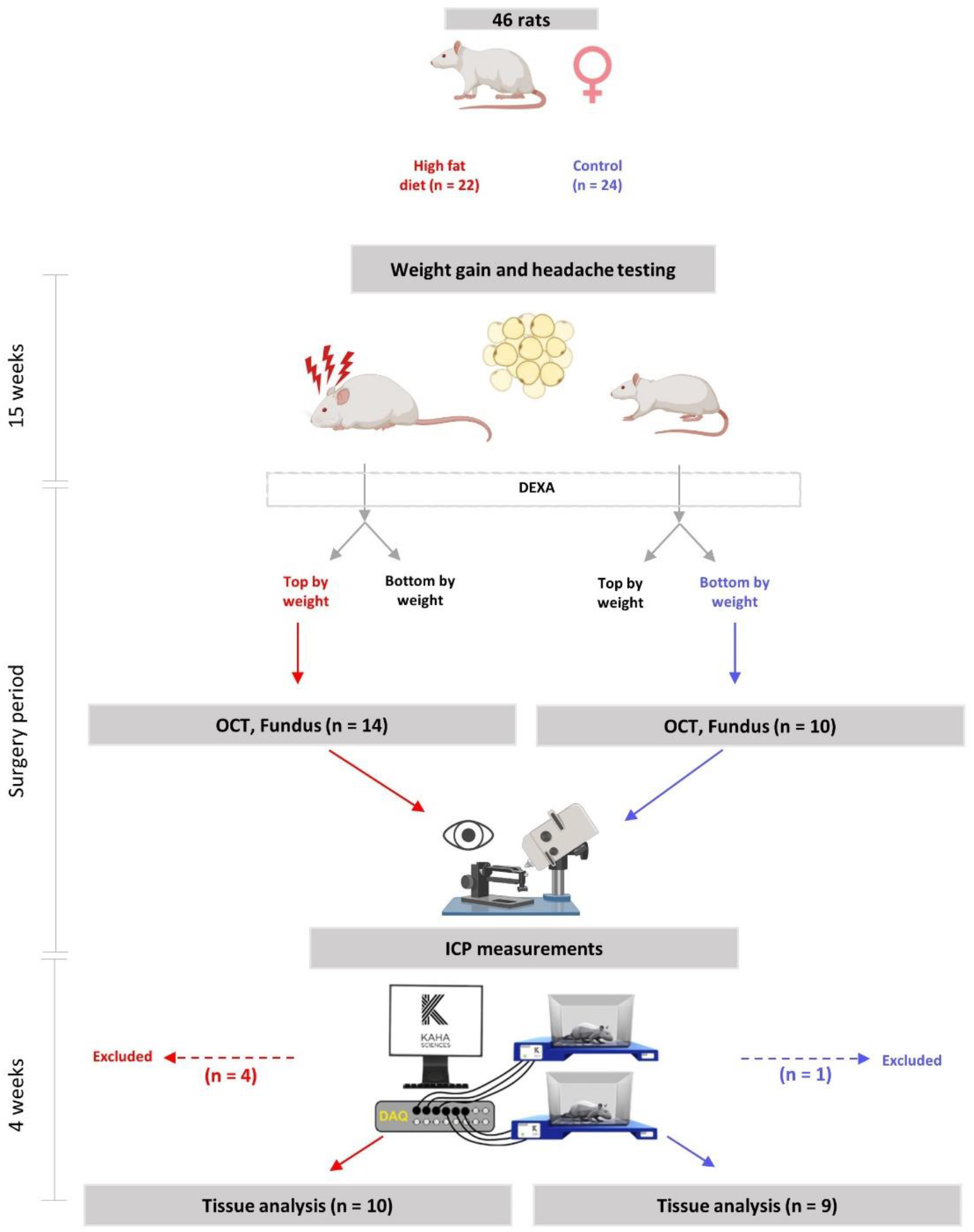
Schematic diagram of experiment. Schematic diagram detailing the experimental paradigm and usage of animal during the study. Female rats were on high fat diet or control matched diet for 15 weeks. Von Frey testing was performed during the whole diet period. Following the 15 weeks, Von Frey testing, DEXA scanning and all the eye examinations were performed followed by implantation with ICP telemetric probe for continuous ICP recording (day 0). During the recovery period (day 0-8), the rats were kept single-housed to recover from the surgery and then co-housed (from approximately day 8). Day 8-30 the rats were not affected by anesthesia or any drugs and ICP recordings was continued until day 30.

### DEXA scanning

Body composition was estimated using a Lunar PIXImus DEXA scanner (GE Medical systems, Chicago). Scans were analyzed using LUNAR PIXImus 2 software in the region of interest generating values for lean mass (LM), fat mass (FM) and bone mass (BM). The region of interest was set to S1/L6 vertebral space to the L1/T12 vertebral space, thus scanning the abdomen ^48^.

### ICP telemeter implantation and ICP recording

Selected animals from the DIO and control groups were implanted with ICP probes for continuous recordings (n=23), which was carried out as previously described ^18^. In brief, rats were anaesthetized with hypnorm/midazolam or ketamine/xylazine and the telemeter body was implanted in the abdominal region. The ICP catheter was tunneled under the skin to the parietal bone where the sensor tip was placed and secured epidurally. After surgery, the rats were placed in their home cages upon a Smartpad and ICP recording started. The telemeter sampled ICP at 2 kHz, where the Smartpad lowpass filter at 1 kHz gave a final sampling frequency of 1 kHz. After surgery and the two following days the rats received post-surgery treatments and were allowed to recover up to 7 days as described earlier ^18^. On the postoperative days some of the rats lost weight (<15%) and from day 7 weight increased as observed with this kind of procedure ^18,19^. At the end of the experiments all animals were euthanized with an overdose of sodium pentobarbital. Some animals had to be excluded due to adverse events such as opening of stiches and injection site reaction.

### ICP analysis

ICP was measured for 30 days after implantation surgery. Presented means are for day of surgery, day 0 (n=9 control, n=10 DIO), days 3, 7 and whole period (n=8 control, n=10 DIO). More specific spectral analysis of the ICP waveform was carried out on a 5-minute stable section of ICP, i.e. without movement artefacts as previously described ^18^.

### Measurement of hind paw and periorbital thresholds after mechanical stimulation

Von Frey testing was performed from diet start until day 0 (Fig.6). The sensitivity of the frontal region of the head (V1 ophthalmic trigeminal dermatome) and hind paw to static mechanical stimulation was measured using an electronic von Frey device fitted with a rigid plastic tip (IITC LifeScience Inc, USA). Testing was carried out as previously described ^49^. The investigator was blinded to the experimental group during the experiment.

### Measurement of light sensitivity

Photophobia was assessed by quantifying time spend in the light during a light/dark box test as previously described ^50,51^. The light/dark box was placed in a dedicated room with video recording where the rats were left to acclimatize before the test (HFD n= 22, CD n=24). Light intensity was set to 500 lux for 20 minutes. Time spend in the light (s) was assessed by blinded observers and defined as the eyes of the rats being visible in the light compartment of the box i.e. exposure to light. Photophobia was tested during their active period and for every round the rats were tested at the same day.

### OCT, ODP image capture and analysis

OCT and ODP of both eyes were performed prior to implantation of the ICP probe on day 0 (DIO n= 14, CD n=10, 48 eyes). OCT and ODP images were captured with a combined digital fundus microscope and image-guided spectral domain OCT system (Phoenix MICRON IV In Vivo Imaging Microscope and OCT2 system, Phoenix Technology). The corneal surface of eyes was topically anesthetized with Phenylephrine Hydrochloride 10% and pupils were dilated with Tropicamide 1%. Guided by the fundus microscope the ONH was centered for the best OCT image. The OCT image was then aligned horizontally, and optimized for best image quality adjusting the polarization, contrast, gamma and dispersion. Peripapillary circular B-scans (1024 A-scans with a radius of 515 μm) were performed with 3-10 image averaging to reduce image noise. Finally, an ODP was captured with the optic disc centered and in focus.

From each eye, the best quality OCT images with a minimum of movement artefacts were analyzed using semi-automatic retinal segmentation software (InSight, Phoenix Technology). Three layers (inner limiting membrane (ILM), plexiform layer (IPL) and Bruch’s membrane (BM)) were automatically segmented and manually fitted for best accuracy, and additionally two layers (retinal nerve fiber layer (RNFL) and inner nuclear layer (INL)) were manually segmented. The thickness of the total retina (TR), the RNFL, the ganglion cell complex (GCC, which consists of the RNFL, the ganglion cell layer (GCL) and the IPL), the IRL and the outer retinal layers (ORL) were extracted using a spreadsheet software. All OCT segmentations and ODPs were blinded for a neuro-ophthalmologist for evaluation. The ODP images were evaluated for obvious pathology including optic disc obscuration, peripapillary halo, vessel engorgement/torsion/obscurations, hemorrhages and choroidal folds. Intraocular pressure (IOP) was measured in non-anaesthetized rats on day 0 using a TONOLAB tonometer (ICARE, Finland). The investigator was blinded to both the experimental group and identity of the rat.

### Histology of eyes

At the end of the study (day 30), eyes were collected and fixed in 4% paraformaldehyde (PFA) in phosphate buffered saline (PBS) (HFD n= 9, CD n=7, 32 eyes). The tissues were cryoprotected by being immersed in raising concentration of sucrose in Sörensen’s phosphate buffer, followed by embedding in gelatin medium and cryosectioned at 12μm (Leica Microsystems GmbH). For evaluation of morphology, sections were stained were hematoxylin-eosin (HTX-Eosin). For measurement analysis, the best quality section from each animal covering the central portion of the ONH, and presenting the retinal nerve fibers entering the optic cup, were used for measuring the retinal nerve fiber bundle (RNFB) thickness at the level of the outer plexiform layer on each side of the ONH as previously described^52^.

### mRNA expression

CP and TG were removed from rats following sacrifice, snap frozen and stored at −80°C prior to RNA extraction. Total RNA was extracted using Trizol reagent (Invitrogen™, Waltham, MA, USA). RNA was converted to cDNA using the Applied Biosystems High Capacity cDNA Reverse Transcription Kit according to manufactures instructions. RT-qPCR was performed using the Applied Biosystems QuantStudio 6 Pro. Taqman Gene Expression Assays (Life Technologies) were used to assess gene expression. Reactions were carried out in 384 well plates, single plex 10μl reaction volumes, using Taqman Gene Expression Master Mix (Applied Biosystems) where samples were ran in triplicate. Genes expression was assessed using Taqman primer/probes sets, the genes assessed were *Aqp1* (Rn_00562834_m1), *Slc12a2* (Rn_00582505_m1), *Calca* (Rn_01511353_g1), *Ramp1* (Rn_01427056_m1), *Trpv1* (Rn_00583117_m1), *Tnf* (Rn_99999017_m1) and *Ilb1* (Rn_00580432_m1). Expression of the target gene was normalized to the expression of *Actb* (Rn_00667869_m1) for TG and *Gapdh* (Rn_01775763_g1) for CP. The relative expression of the gene is presented as fold change (2^-((ΔCt subject)-(meanΔCt control)), statistics are done on ΔCt values and arbitrary units.

### Western Blot

Protein was obtained after extracting RNA from the samples with Trizol reagent (Invitrogen™, Waltham, MA, USA). Protein was extracted from the same sample, precipitated and washed accordingly to the manufactures protocol. The protein was solubilized in lysis buffer with 1% sodium dodecyl sulfate. β-mercaptoethanol was added as a reducing agent making up 5% of the solution. Protein concentration was determined in triplicates on NanoDrop Spectrophotometer (NanoDrop™ 2000c, Thermo Scientific™, USA). The proteins were separated using 4-10 % bis-tris gel (Invitrogen™, USA) under gel electrophoresis and transferred to a polyvinylidene difluoride membrane (iBlot™, USA). The membrane was blocked and incubated in primary antibody overnight diluted 1:1000 for Aqp1 (ab168387, Abcam), Lamin-B1 (ab65986, Abcam) and Nkcc1 (14581, Cell Signaling). Membranes were incubated in secondary antibody diluted 1:10000 and the protein bands were detected using chemiluminescent (Amersham™ ECL™, USA). The reaction was captured on luminescent image analyzer (LAS-4000 Luminescent Image Analyzer, Fujifilm, Japan). Protein band was quantification using ImageJ (NIH, USA). The relative protein expression is related to the loading control.

### Blood glucose

Blood glucose was measured on day 30, where the rats had been fasted overnight for at least 14h. Blood from the lateral tail vein was used to asses blood glucose (mmol/L) using glucometer ACCU-CHEK Aviva (Roche Diagnostics, Denmark).

### Statistics

Statistical analysis was performed using Graphpad prism 9 (Graphpad Software Inc, USA). Data presented as mean±SEM unless otherwise stated. Data normality assessed via Shapiro-Wilk normality test. Where data was normally distributed unpaired two-tailed t-tests (equal variance) or Welch’s test (unequal variance) was employed, whereas non-parametric data was assessed via Mann-Whitney U. Pearson’s correlation coefficient (r) was used for assessing correlations. Results were judged significant at P<0.05.

## Acknowledgements

Professor Niklas Rye Jørgensen and PhD Maria Ellegaard Larsen at Glostrup Research Institute for the use of the DEXA scanner.

Professor Birgitte Holst, Professor Nanna MacAulay and ass professor Jonathan Wardman at Copenhagen University for valuable discussions.

## Notes

### Competing Interest Statement

The authors have declared no competing interest.

